# Genome-wide Segregation of Single Nucleotide and Structural Variants into Single Cancer Cells

**DOI:** 10.1101/117663

**Authors:** John Easton, Veronica Gonzalez Pena, Donald Yergeau, Xiaotu Ma, Charles Gawad

**Author notes:** equal contribution (John Easton, Veronica Gonzalez Pena). Correspondence: Charles Gawad.

## Abstract

We present a new approach for determining comprehensive variant profiles of single cells using a microfluidic amplicon-based strategy. This method can be used to reconstruct the clonal architecture and mutational history of a malignancy using all classes and sizes of single nucleotide and structural variants, providing insights into the temporal changes in mutational classes and processes that led to the development of a cancer. Using this approach, we interrogated single cells from a patient with leukemia, determining that processes producing structural variation preceded single nucleotides changes in the development of that malignancy.

## Introduction

Recent technological advancements have enabled the sequencing of genomes of single cells, a technologically challenging process that starts with a single molecule of DNA[1]. By bringing genomics to the cellular level, we have begun to segregate mutations to distinct cellular populations, enabling us to define the population genetic diversity and clonal structures of complex tissues. Initial studies have provided unexpected insights into the contributions of somatic mosaicism to human development and disease[2, 3]. This has been especially true in cancer where intratumor genetic heterogeneity has provided the opportunity to trace back the mutations and mutational processes that resulted in the formation of a malignancy[4, 5].

Contemporary strategies for amplifying and interrogating the genomes of single cells have resulted in the ability to segregate single nucleotide or copy number variants (CNV) into single cells. However, due to the tradeoffs in genome coverage and uniformity using current amplification strategies, we have had limited success comprehensively detecting both variant classes in the same cells[6]. In addition, most strategies for detecting CNV rely on differences in read depth at specific locations relative to a reference, which are only able to detect large regions of alteration[7]. Further, read depth does not provide information on other classes of structural variation (SV), including: translocations, inversions, insertions of novel sequence, or interspersed copy number gains. Isothermal methods that provide sufficient breadth of genomic coverage to identify most single nucleotide variants (SNV) do not have adequate uniformity in coverage depth to acquire CNV information[5]. Quantitative PCR-based methods have been developed to detect both CNV and SNV in the same cell, but they are only able to interrogate a small number of variants due to limitations in multiplexing, which hampers the investigator's ability to accurately determine the clonal structures[8, 9].

In the present study, we report a strategy for segregating hundreds of any variant type to individual cells in an accurate, cost-effective, and efficient manner. We accomplish this by using multiple displacement amplification for maximal genome recovery of each cell[10], followed by detecting all classes of structural variation using the breakpoint sequence at the site of rearrangements rather than measuring read depth[11]. We first perform whole genome sequencing to comprehensively characterize all types of somatic variation in a bulk population (Figure 1A). We then isolate and amplify the genomes of single cells using a microfluidic platform, followed by microfluidic high-throughput amplicon-based confirmation of bulk variants and determination of single cell mutation profiles (Figure 1A). We then use the single cell mutation profile to infer the clonal structure and mutational history of that malignancy[5, 12].

**Figure 1.**
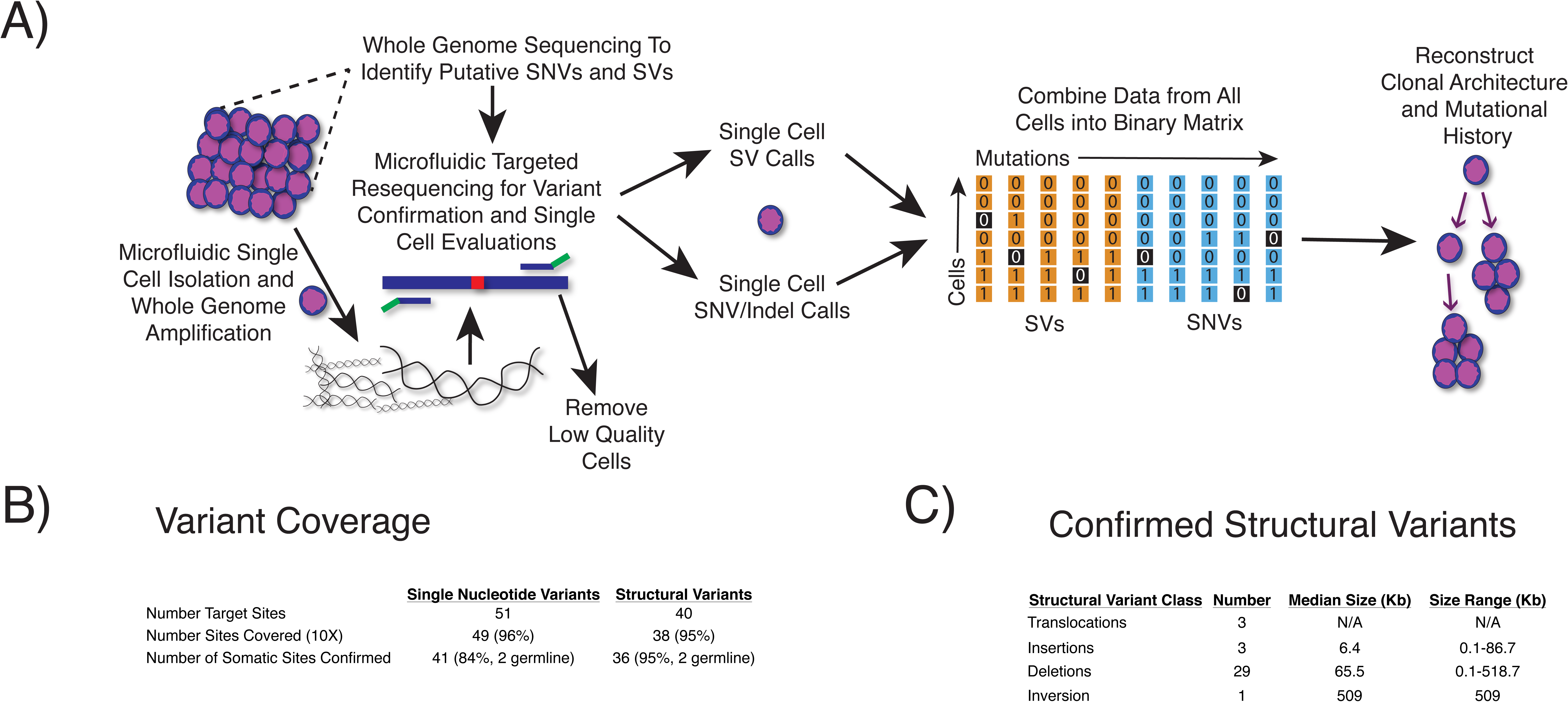
Overview of Experimental Protocol and Performance A) Whole genome sequencing is first performed to determine the comprehensive mutation profile of the sample, followed by variant confirmation using targeted resequencing. Single cells are then isolated, followed by amplification of the variant sites, variant calling, and binary matrix construction to determine the clonal structure. B) Overview of putative variant site coverage and confirmation rates. C) Class and size of confirmed structural variants.

## Results

As an example, we used this approach to segregate mutations into single cells from samples taken from a child that was diagnosed with acute lymphoblastic leukemia. We first performed whole genome, exome, and RNA sequencing to characterize the somatic variants in that sample as part of the Pediatric Cancer Genome Project[13, 14], and were able to identify 51 putative SNVs and 38 SVs (Figure 1B). We then performed microfluidic amiplicon-based resequencing and were able to cover 96% of the 91 target sites at 10X coverage depth in bulk samples. We confirmed 36 somatic SVs that included 3 translocations and an inversion, as well as 3 insertions and 29 deletions that ranged in size from 0.1 to 518.7 Kb (Figure 1C, Table S1). In addition, we confirmed 41 SNVs that had a strong enrichment for C to T transitions and C to G transversions within an APOBEC motif, in agreement with previous studies of this type of leukemia (Table S2)[5, 9].

We then captured 168 cells in two microfluidic chips where 128 were confirmed to be single cells by microscopy. As shown in figure 2, after creating a binary matrix of all cells and mutation calls, we then retraced the mutational history of the tumor by grouping cells and mutations into clusters using probabilistic-based clustering, which revealed two distinct clonal populations that were formed from 3 distinct mutational clusters. There was one cluster of poor performing assays (red cluster, 6.3% of assays) that contributed to the overall estimated false negative rate (allelic dropout + 2 * locus dropout) of 23.8%, as well as a small cluster of double cells despite putative identification of single cells by microscopy (red cluster, 9.6% of cells). Closer examination of the mutations in the clusters showed a shared ancestral cluster that had only acquired SV, followed by separate mutation clusters of SNVs and a smaller number of SVs being acquired as the two clones evolved (Figure 2).

**Figure 2.**
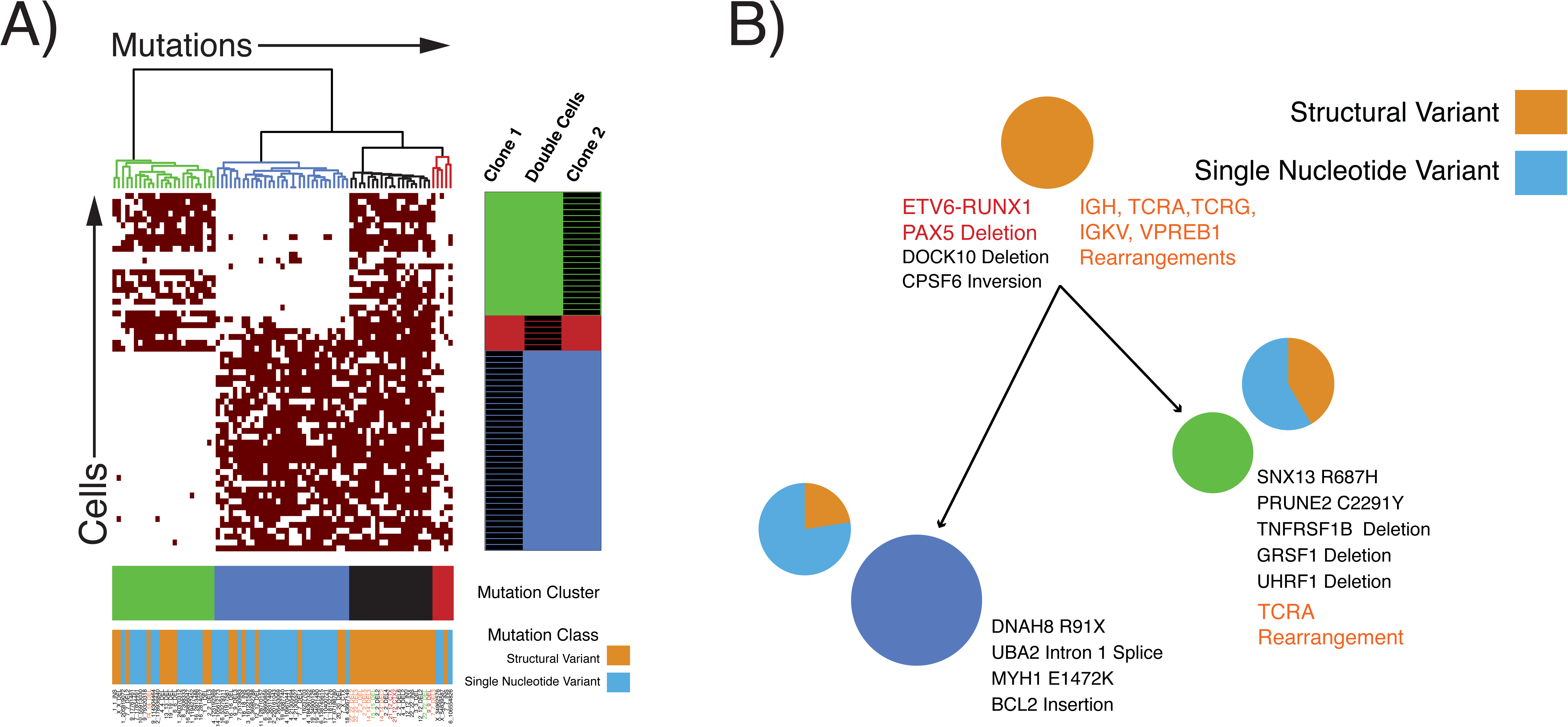
Childhood Leukemia Example A) Heatmap depicting clustering patterns of cells and mutations. Three clear mutation clusters segregate into two distinct clonal populations. The ancestral shared mutation cluster is composed entirely of structural variants. B) Minimal spanning tree showing the relative size of and genetic distance between each clone. Known driver mutations are depicted in red, immune receptor rearrangements in orange, and putative drivers of clonal expansions in black. The relative contribution of single nucleotide and structural variation to each clone are also represented by the pie charts adjacent to each clone.

These data further support the assertion that there was a distinct process creating SV that drove tumor evolution and preceded the process that induced the later SNVs. In addition, it has been shown that SV in this type of leukemia have a signature of immunoglobulin recombination activating genes (RAG1/2) while the SNVs have an APOBEC signature. Consistent with this, the ancestral clone that only harbored SV had rearrangements of most of the RAG target immune receptor genes. However, that variation was insufficient to produce malignant transformation, which required the APOBEC-mediated SNVs that drove later evolution of the leukemic cell genomes. Thus, we are not only able to order the sequences of genetic events, but also the underlying mutational processes that drove malignant transformation of the disease as those normal fetal hematopoietic precursors evolved into leukemia over several years[15].

## Discussion

Single-cell sequencing is a powerful tool for deconvoluting the mutational histories of tumors. Both single nucleotide and structural variants can contribute to malignant transformation, and they frequently occur as a result of distinct mutational processes. By acquiring information on all subtypes and sizes of both variant types in the same cells, we have provided a strategy for learning the order in which these events occur, and potentially, how they cooperate in the development of human cancer. This strategy can be applied to deconvolute the mutational histories of all cancer types as we try to understand the processes that drive the transformation of normal cells into cancer.

## Methods

### Samples and Bulk Sequencing

Samples from this patient were obtained following approval of the St. Jude Children's Hospital Institutional Review Board and informed consent of the family. Mononuclear cells were isolated from fresh bone marrow samples using Ficoll-Paque (GE Life Sciences), followed by standard cryopreservation. Whole genome, exome, and RNA-sequencing were performed on DNA isolated from the leukemia cells as part of Pediatric Cancer Genome Project[13]. The methods for producing and analyzing the data to generate the putative variant lists have been previously published[14]. We attempted to validate all SVs, as well as SNVs that resided in coding regions.

### Single-Cell Isolation and Whole Genome Amplification

Vials of leukemia cells were thawed using the ThawSTAR System (Biocision), followed by one 15ml wash in prewarmed RPMI with 1% bovine serum albumin (Sigma). The cells were washed four additional times using Fluidigm wash buffer according to the manufacturer. Cells were filtered using a 15uM strainer (PluriSelect), followed by counting and viability estimation using Luna-FL counter (Logos Biosystems). The cells were then resuspended at a final concentration of 300 cells/ul before mixing with suspension reagent at a ratio of 4ul of cells to 6ul of suspension reagent. The cells were then loaded into a small Fluidigm C1 DNA sequencing microfluidic chip, followed by whole genome amplification according to the manufacturer's instructions.

### Targeted Sequencing of Variants in Single Cells

Single cell whole genome amplification products underwent targeted sequencing using the Access Array System according to the manufacturer's instructions as previously reported (Fluidigm)[5]. Primers were designed across SNV sites or SV breakpoints using BatchPrimer3 (Table S3)[16] adding common sequences according to the Access Array instructions (Fluidigm). The sequences of the primers are listed in supplementary table 1. Bulk DNA from both tumor and a remission sample as a germline control were run on each Access Array chip to confirm bulk variants, as well as to insure successful amplification on the chip. Barcoded amplicons from four Access Array chips were pooled and run on a MiSeq using 2X150bp reads (Illumina).

### Data Processing, Variant Confirmation, and Clonal Structure Determination

Trimming, alignment, and SNV calling were performed as previously reported[5]. SVs were first confirmed in the bulk sample by determining the number of reads that had an exact match to the 30bp sequence that spanned the breakpoint. Variants were considered present if greater than 40 reads were an exact match of the 30bp that spanned the breakpoint. That threshold was chosen to minimize false positive calls due to sample cross contamination or incorrect demultiplexing of sequencing reads. Chambers that were visually confirmed to contain one cell and samples that covered at least 80% of target SNV sites at a depth of 10X or greater were retained for further analyses. SV calls were then made in each single cell, which were combined with SNV calls to create a binary matrix containing all cells. The number of clones were estimated after performing hierarchical clustering. Cells were then assigned to clusters using an expectation-maximization algorithm, followed by minimal spanning tree construction as previously reported[5, 12].

## Acknowledgements

The authors would like to acknowledge Jinghui Zhang and Jim Downing for their constructive advice on the project. In addition, the authors would like to thank the members of the Pediatric Cancer Genome Project, especially Charles Mullighan and Ching-Hon Pui who lead the Hematological Malignancies Program. All authors are supported by ALSAC. C.G. is also supported by the Burroughs Wellcome Fund, Leukemia and Lymphoma Society, Hyundai Hope on Wheels, and the American Society of Hematology.

The data are available in the short read archive accession ID.

## Author contributions

: C.G., J.E., V.G. designed research; V.G., D.Y. performed research; C.G. contributed new reagents/analytic tools; C.G., X.M., and J.E. analyzed data; and C.G., V.G., and J.E. wrote the paper.

---

**Table S1.** List of Confirmed Structural Variants

**Table S2.** List of Confirmed Single-Nucleotide Variants

**Table S3.** Primers Used for Targeted Resequencing of Variant

